# Plant endophytic bacteria: a potential resource pool of electroactive microorganisms

**DOI:** 10.1101/2020.10.11.334912

**Authors:** Lijun Ling, Zibin Li, Caiyun Yang, Shenglai Feng, Yunhua Zhao, Wenxia Ma, Lu Lu

**Author notes:** Address correspondence to Lijun Ling,. 967 Anning East Road, Anning District, Lanzhou City, Gansu Province, China (Lijun Ling).

## Abstract

Electroactive microorganisms play a significant role in microbial fuel cells (MFCs). These devices, which are based on a wide microbial diversity, can convert a large array of organic matter components into sustainable and renewable energy. At present, electricity-producing microorganisms are mostly isolated from sewage, anaerobic sediments and soil, however, the sources are very limited. For a more comprehensive understanding of the electron transfer mechanism of the electricity-producing microorganisms and the interaction with the environment, it is necessary to obtain a thorough understanding of their resource distribution and discover potential resources. In this study, plant tissues were selected to isolate endophytic bacteria, and the electrochemical activity potential of those bacteria was evaluated by high-throughput screening with a WO3 nanoprobe. Twenty-six strains of endophytic bacteria were isolated from plant tissues belonging to Angelica and Sweet Potato, of which 17 strains from 6 genera had electrochemical activity, including Bacillus sp., Pleomorphomonas sp., Rahnella sp., Shinella sp., Paenibacillus sp. and Staphylococcus sp.. Moreover, the electricity-producing microorganisms in the plant tissue are enriched. Microbial community analysis by high-throughput sequence indicated that Pseudomonas and Clostridioides are the dominant genera of MFC anode inoculated with angelica tissue.Staphylococcus and Lachnoclostridium 5 are the dominant genera in MFC anode inoculated with sweet potato tissue. And the most representative Gram-positive strain Staphylococcus succinus subsp. succinus H6 and plant tissue-inoculated MFC were further analyzed for electrochemical activity. After nearly 1500 h of voltage monitoring and cyclic voltammetry analysis, the results showed that a strain numbered H6 and plant tissue-inoculated MFC had a good electrogenerating activity.

**Importance:** Some biological characteristics of microorganisms are inextricably linked to their living environment. For plant endophytes, some of their biological characteristics have a profound impact on the host. The discovery of the production of electrobacteria in plants helps us to understand the interaction between microorganisms and plants more deeply. For example, there may be intercellular electron transfer between the internally producing bacteria and nitrogen-fixing bacteria to improve the efficiency of nitrogen fixation. In addition, there may be a connection between the weak electrical signal of the plant and the the endophytic electricity-producing microorganismsThe discovery of electricity-producing bacteria in plants also brings a more comprehensive understanding of the distribution of electricity-producing microbial resources and the mechanism of origin.

## Introduction

Microbial fuel cells (MFCs) are fuel cells that convert chemical energy to electrical energy, using microbes as catalysts^(1)^. Because of the environmental characteristics, MFCs have been widely used as new energy production devices. In addition, MFCs can significantly improve sewage treatment efficiency. The substantial increase in power density has become a bottleneck for extensive applications of MFCs^(2)^. After more than one hundred years of development, the performance of MFCs has been greatly improved, and the power density reaches 2.08 kWm^-3(3).^ Many studies on optimizing the design and construction of MFCs were undertaken to solve the problem, while the importance of electroactive microorganisms and electricity generation mechanisms were often poorly emphasized. Electroactive microorganisms, which possess the ability to transfer electrons to extracellular electron acceptors in their respiration, have an important effect on the performance of MFCs ^(4).^ Microbial power generation had been discovered at least one hundred years ago by Porret et al. ^(5)^. Besides the electroactive microorganisms had been found in diverse environments ^(6).^ These bacteria play an important role in bioelectrochemistry, geochemical cycles, biocorrosion, and environmental remediation ^(7, 8)^.

Since one of the most promising applications of MFC is the treatment of wastewater, most of the application research of MFC is directed at domestic sewage treatment plant wastewater, paper mill wastewater(9, 10), etc. So, separation and screening of electricity-producing microorganisms are also concentrated in these environments among. To date, most of the electricity-producing microorganisms found are Gram-negative bacteria. The extracellular electron transfer mechanism of electricity-producing microorganisms is also mostly directed against Gram-positive bacteria. The limited Gram-positive electricity-producing microorganisms prevent us from understanding the electricity-producing microorganisms more comprehensively and in-depth. Therefore, it is particularly important to discover new electricity-producing microbial resources.

The present study aimed to gain knowledge on the diversity, uniqueness, and biogeography of electroactive microorganisms inhabiting the environment of plants. For this purpose, different microbial samples that originated from plant underground rhizomes were studied using cultivation-based approaches. Isolates were characterized through phylogenetic analysis of 16S rRNA gene sequences, and the power production performance of the representative strains and Plant tissues were investigated. At the same time, inoculate the plant tissue with electricity-producing microorganisms into MFC, and after continuous and stable operation for many cycles, high-throughput analysis of anode community structure. In this paper, the feasibility of plant endophytic bacteria as resources of electroactive microorganisms was revealed, and a new resource library of screening the resources of electroactive microorganisms was found.

## Material and Methods

### Isolation and identification of endophytes in plants

The underground roots and stems of two kinds of plants, sweet potato (*Dioscorea esculenta* Burkill) and angelica (*Angelica sinensis* Diels), were collectioned from Meichuan town, Min County, CN (PR-MC/RS-MC, 34°43’N, 104°03’E). The complete roots of sweet potato and stems of angelica were surface sterilized with 70% ethanol for 1 min, followed by 5% sodium hypochlorite for 5 min, and then rinsed five times in sterile deionized water (dH_2_O). The effectiveness of surface sterilization was verified by spreading the final rinse water onto NA (nutrient agar). Sterilized tissue was slit, the internal tissue was homogenized at a sterile workbench, and 100 μL of the homogenate was added to NA plates. The plates were incubated at 37 °C until visible colonies appeared. Colonies of varying morphologies were selected and purified by cross streaking onto NA plates. Purified strains were suspended in NA broth supplemented with 10% (v/v) glycerol and maintained at −80°C ^(11).^ Electrochemically active bacteria were distinguished by a high-throughput method using a WO_3_ Nanocluster Probe^(12)^. First, 96-well plates with identical transparency for each well were used for high-throughput evaluation of microorganisms. A mixture of 120 mL of a bacterial solution in medium and 80 mL of a 5% sterile WO_3_ nanocluster-containing suspension was added to 96-well plates, followed by the immediate addition of 80 mL of paraffin oil to ensure anaerobic conditions. The plate was incubated at 37°C, and color development was checked after 12 h. Triplicates were cultivated in parallel. the strains whose color has changed were separated and preserved, and then the electrochromic properties were identified again.

DNA of pure cultures was prepared according to established protocols, with modifications ^(13).^ Briefly, cultures were grown overnight in 10 mL of NA broth at 37 °C, and cells were collected by centrifuging at 12 000 × g for 1 min, followed by washing in ddH_2_O. Cells were lysed by vortexing and adding glass beads. Lysates were clarified, and cDNA was extracted using the phenol/chloroform/isoamyl alcohol method. Extracted DNA was precipitated in an equal volume of isopropanol at room temperature. DNA was resuspended in TE buffer and stored at −20 °C.

The 16S rRNA sequence of the large ribosomal ribonucleic acid subunit (LSU rRNA) gene region was PCR amplified using the 27F and 1492R primers. PCRs contained 5 ng of DNA template, 0.2 mM 27F (5′ AGAGTTTGATCCTGGCTCAG 3′) and 1492R primers (5′ GGTTACCTTGTTACGACTT 3′), 0.4 U of Taq polymerase and 3.7 mM MgCl_2_ (Takara) in PCR buffer (total volume, 50 µL). The PCR conditions were as follows: 5 min denaturation at 94 °C; 30 cycles at 94 °C for 30 s, 54 °C for 30 s, and 72 °C for 50 s; followed by a final 72 °C extension for 10 min. PCR products were verified on 1% agarose gels.

PCR products were sequenced by a gene sequencing company (TianQi Gene, China). Sequences were compared by BLAST searches ^(14)^ and aligned to related sequences retrieved from GenBank using the multiple alignment program. Nucleotide sequences obtained in this study were submitted to GenBank (http://www.ncbi.nlm.nih.gov/BLAST/).

### Microbial community analysis

Immediately after the reaction, the carbon felt anode was taken out of the reactor, and an appropriate amount of carbon felt was cut at different positions of the carbon brush with sterilized scissors. The cut carbon felt is gently rinsed in sterilized deionized water to remove non-electrode microorganisms or impurities attached to it. Shred the fiber with scissors. Simultaneous collection of anolyte for biodiversity analysis of electricity generation.

The genomic DNA was extracted from the samples using the MOBIO PowerSoil DNA Isolation Kit. The obtained DNA was detected by agarose electrophoresis, and the concentration of the extracted genomic DNA was determined by Nanodrop 2000.

After genomic DNA was extracted from the sample, the V3 + V4 region of 16S rDNA was amplified using specific primers with barcode. The primer sequences were: 341F: CCTACGGGNGGCWGCAG; 806R: GGACTACHVGGGTATCTAAT^(15)^. The PCR amplification products were cut and recovered, and quantified using a QuantiFluorTM fluorometer. The purified amplified products were mixed in equal amounts, connected to a sequencing adapter, and a sequencing library was constructed. The PCR products were sequenced on an Illumina Miseq platform (GENEDENOVO Biotechnology Co., Ltd., Guangzhou, China). The sequencing data of bacteria was analyzed using QIIME software package and UPARSE pipeline according to the standard protocols. The operational taxonomic units (OTUs) at 97% similarity were used to cluster the sequences^(16)^.

### Identification and electrochemical activity analysis of representative strains

#### Strain identification

To further verify the electrogenic performance of endophytic bacteria, a representative strain was selected for electrochemical performance analyses.

The colony morphology of isolates was inspected on NA medium by direct observations or the use of a stereomicroscope. Conventional biochemical tests, such as KIA, oxidase, catalase, Gram staining and indole tests, were performed according to Bergey’s Manual of Systematic Bacteriology, volumes 1 and 2^(17, 18)^. Physiological tests were also performed in LB medium to determine the physiological properties of the isolates.

Phylogenetic trees were reconstructed from evolutionary distance data on Kimura’s two-parameter correction ^(19)^ using the neighbor-joining method ^(20)^ and MEGA software version 7.0. Confidence levels of the clades were estimated from bootstrap analysis (1000 replicates) ^(21)^.

#### MFCs construction

Cubic-type and bottle-type MFCs described previously ^(22)^ were used in this study. The Cubic-type MFCs were used to detect the electrochemical activity of the strain. Strains were cultured in LB medium Strains were cultured in LB medium until the OD_600_ was 1.5. Cells (5%) were inoculated into 13 ml anaerobic plexiglass chambers that contained only a carbon felt electrode (7 cm^2^) as an electron acceptor. The medium in the chambers was identical to that used for growth. The cathode chamber was filled with 13 mL of 50 mM potassium ferricyanide and 100 mM phosphate buffer (6.8 g/L KH_2_PO_4_ and 17.9 g/L Na_2_HPO_4_·12H _2_O) as an electron acceptor. The anode in the anode chamber was connected via a 1 000-resistor to an identical electrode (cathode) in a cathode chamber. The MFC was conducted at 30°C.

Bottle-type MFCs were used to detect the electrochemical activity of endophyte in plant tissues. Disinfect plant tissue using the method described previously. Sterilized tissue was slit, the internal tissue was homogenized at a sterile workbench, and 3g of the homogenate was added to MFC anode chamber. Both anode and cathode chambers have a volume of 100ml. The projected area of the anode and cathode electrodes are 14 cm^2^. Others are the same as the above reactor.

An MFC without an inoculum was used as a control. The anode solution was replaced at the same time after the voltage dropped to 40 mV. The two chambers were separated by a proton exchange membrane (DuPontTM Nafion^®^ NR-212), as previously described. All parts of the reactors were submerged in 75% ethanol for 1 day before use. The culture medium was sterilized at 121°C for 20 min by autoclaving.

#### Electrochemical analysis

Cell voltage was automatically monitored by a data acquisition system (USB3202, ART, China) every 2 hours during electricity generation. Polarization data were taken after 15 min at each external resistance at the beginning of a single operation cycle^(23)^. The power density was calculated from P=IU/A, where I is the current, U is the voltage (both were recorded by the potentiostat), and A is the area of the anode electrode of the MFC. Internal resistance, R_int_, was measured by the power density peak. Each treatment group was set up in three parallel.

Cyclic voltammetry (CV) (CHI 760E, Chenhua, China) was performed on the supernatant of the centrifuged anode suspension to characterize the electrochemical activity of the microbial metabolites. After a period of a voltage rise period and a voltage stabilization period, the reference electrode to the anode chamber for CV experiments was established. During voltage stabilization, the anode suspension was poured out and centrifuged at 10 000 rpm (∼10 200×g) for 5 min. A carbon electrode (diameter of 3 mm) was used as the working electrode, a platinum electrode was used as the counter electrode, and a saturated calomel electrode (Gaoss Union, China) was used as the reference electrode. The reduction and oxidation reactions were observed in the range of +0.6 V to −0.6 V, respectively. CV was performed in a potassium ferricyanide solution at a scan rate of 5 mV/s.

## Results

### Isolation of endophytic bacteria from underground roots of plants

The isolates revealed 11 strains belonging to 6 genera from angelica, as shown in **Table 1**. Furthermore, the samples from sweet potato yielded a total of 15 strains belonging to 5 genera, indicating that the number of endophytes was higher in sweet potatoes than in angelica, but the level of diversity was lower in sweet potatoes than in angelica.

**Table 1.** Distribution and electrochemical activity of endophytic species

| Strain Serial | GenBank | Generic | Host | Electrochemical |
| --- | --- | --- | --- | --- |
| Code | Login ID |  |  | activity |
| D1 | MK764966 | *Pseudomonas sp.* | Angelica sinensis | - |
| D2 | MK764967 | *Pseudomonas sp.* | Angelica sinensis | - |
| D3 | MK764968 | *Pseudomonas sp.* | Angelica sinensis | - |
| D4 | MK764969 | *Pseudomonas sp.* | Angelica sinensis | - |
| D5 | MK764970 | *Stenotrophomonas sp.* | Angelica sinensis | - |
| D6 | MK764971 | *Streptomyces sp.* | Angelica sinensis | - |
| D7 | MK764972 | *Bacillus sp.* | Angelica sinensis | + |
| D8 | MK764973 | *Pleomorphomonas sp.* | Angelica sinensis | + |
| D9 | MK764974 | *Pleomorphomonas sp.* | Angelica sinensis | + |
| D10 | MK764975 | *Rahnella sp.* | Angelica sinensis | + |
| D11 | MK764976 | *Rahnella sp.* | Angelica sinensis | + |
| D12 | MK764977 | *Shinella sp.* | Angelica sinensis | + |
| H1 | MK764978 | *Bacillus sp.* | Ipomoea batatas<br>(L.) Lam. | - |
| H2 | MK764979 | *Paenibacillus sp.* | Ipomoea batatas<br>(L.) Lam. | + |
| H3 | MK764980 | *Shinella sp.* | Ipomoea batatas<br>(L.) Lam. | - |
| H4 | MK764981 | *Agrobacterium sp.* | Ipomoea batatas<br>(L.) Lam. | - |
| H5 | MK764982 | *Paenibacillus sp.* | Ipomoea batatas<br>(L.) Lam. | + |
| H6 | MK764983 | *Staphylococcus sp.* | Ipomoea batatas<br>(L.) Lam. | + |
| H7 | MK764984 | *Staphylococcus sp.* | Ipomoea batatas<br>(L.) Lam. | + |
| H8 | MK764985 | *Staphylococcus sp.* | Ipomoea batatas<br>(L.) Lam. | + |
| H9 | MK764986 | *Bacillus sp.* | Ipomoea batatas<br>(L.) Lam. | + |
| H10 | MK764987 | *Staphylococcus sp.* | Ipomoea batatas<br>(L.) Lam. | + |
| H11 | MK764988 | *Staphylococcus sp.* | Ipomoea batatas<br>(L.) Lam. | + |
| H12 | MK764989 | *Shinella sp.* | Ipomoea batatas<br>(L.) Lam. | + |
| H13 | MK764990 | *Shinella sp.* | Ipomoea batatas<br>(L.) Lam. | + |
| H14 | MK764991 | *Shinella sp.* | Ipomoea batatas<br>(L.) Lam. | + |

+: positive reaction; -: negative reaction

Electrochemically active bacteria were distinguished by a high-throughput method using a WO_3_ nanocluster probe. After identification, a total of 15 strains of electrochemically active bacteria belonging to 6 genera were obtained. The number of sweet potato samples was the largest, and the isolates were concentrated in *Staphylococcus sp*. In *Angelica sinensis* samples, *Pleomorphomonas sp*., *Rahnella sp*. and *Shinella sp*. showed electrochemical activity, as shown in **Figure 1**. The results showed that the electroactive microorganisms were rich in the plants investigated.

**Figure 1.**
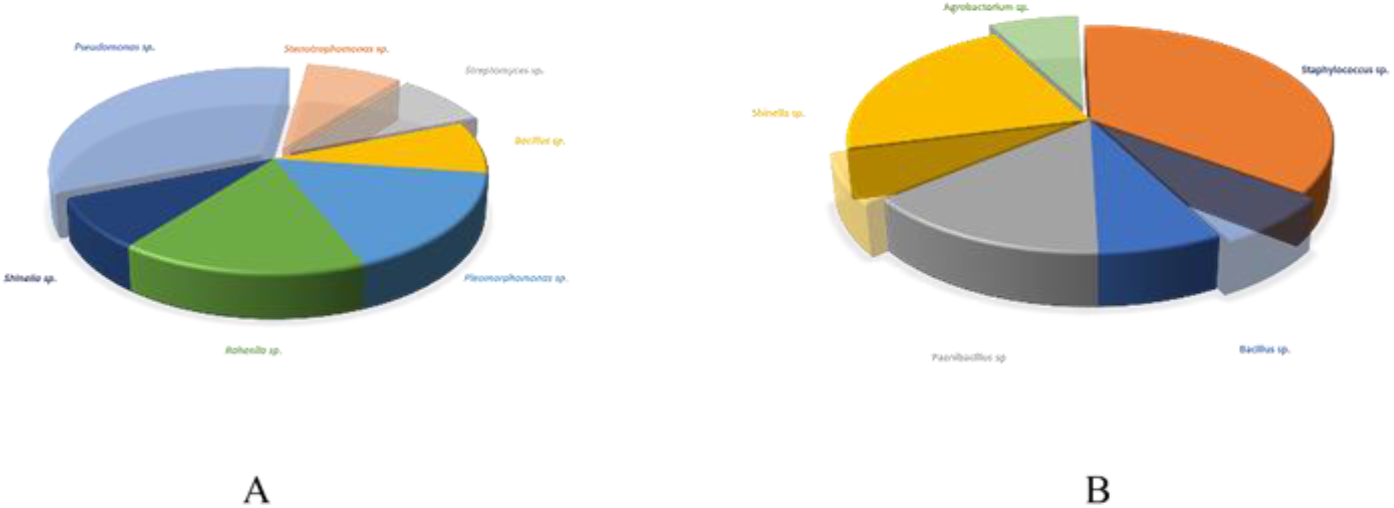
Distribution of endophytic bacteria and electrochemically active bacteria in *Angelica sinensis* (A) and sweet potato (B). The transparent part shows no electrochemical activity.

### Microbial community structure of anode and cathode biofilms

After 5 cycles of MFC inoculation with angelica and sweet potato tuber tissues, the composition and relative abundance of bacterial communities in MFC anolyte and anode electrode membranes were determined at the phylum and genus levels, as shown in Figure 2. The bacteria in the MFC anolyte and anode electrode inoculated with angelica tissue are mainly concentrated in the Proteobacteria, of which account for 75.08% and 68.38% respectively. The microorganisms in the anolyte of MFC inoculated with sweet potatoes are different from Angelica MFC. Most of them belong to the Firmicutes, accounting for 98.14%. The microbial composition of the anode biofilm and anode liquid is also very different. Proteobacteria accounted for 75.34%, and thick-walled bacteria accounted for only 24.59%. Chloroflexi and Bacillus were only found in Angelica inoculated with MFC. In addition, Bacteroidetes, Fusobacteria, Actinobacteria, Spirochaetes, Cyanobacteria were distributed in the four samples. Overall, the Proteobacteria bacteria are the most abundant on the anode electrode membrane of MFC inoculated with angelica and sweet potato. At the genera level, unclassified genera accounted for 47.73%, 61.03%, 70.16%, and 4.94% of AM, AC, SM, and SC samples, respectively. Among the MFC inoculated with Angelica sinensis, Pseudomonas is the dominant genus in AC, accounting for 13.69%. Lachnoclostridium 5, Clostridioides, Clostridium sensu stricto 18, and Bacillus pumilus

**Figure 2.**
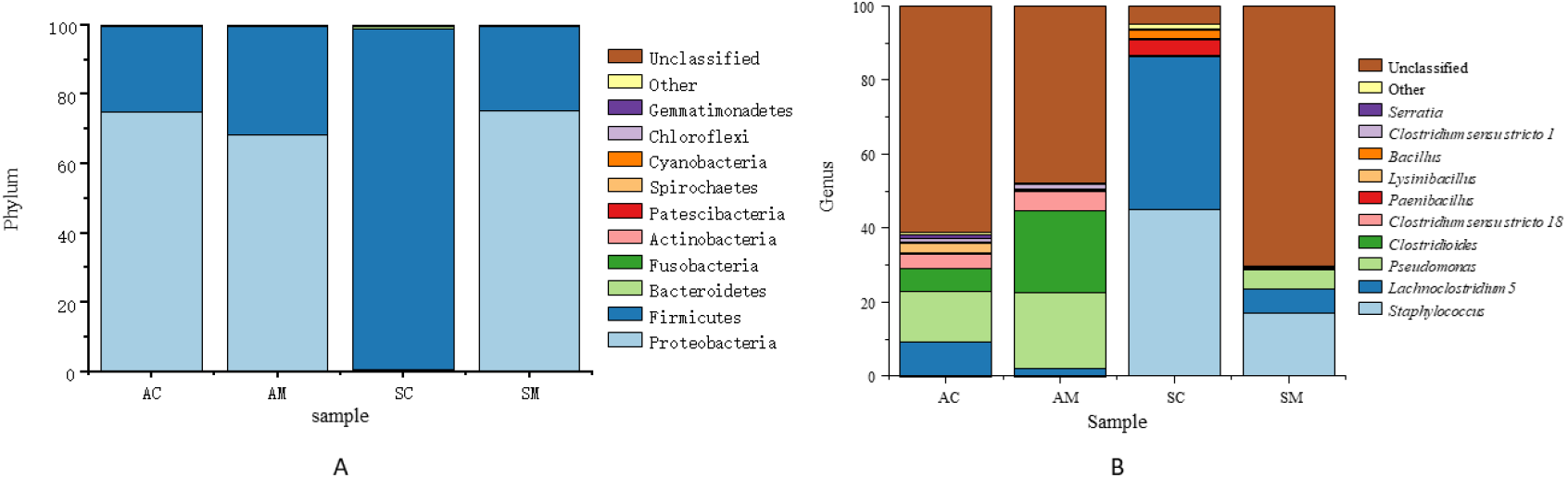
Bacterial classification of phylum level (A) and genus level (B)

The genus *Lysinibacillus* and *Clostridium sensu stricto 1* also occupy a higher proportion, respectively 8.91%, 6.18%, 3.96%, 3.14% and 1.02%. In AM samples, the dominant genera are Clostridioides and Pseudomonas, accounting for 22.04% and 20.72%, respectively. In addition, Clostridium spp., Lachnoclostridium 5 and Clostridium spp. 1 also have a higher proportion, reaching 5.37%, 1.81% and 1.61%, respectively. It is completely different from the MFC inoculated with angelica. Staphylococcus (Staphylococcus) SC and SM are the dominant bacteria in the MFC inoculated with sweet potato, accounting for 45.04% and 17.01%, respectively. The 5 genera of Lachnoclostridium also account for a higher proportion in SC and SM, reaching 41.29% and 6.38%, respectively. Pseudomonas also has certain advantages in SM. In addition, the MFC species inoculated in the two plants have a certain abundance of Bacillus and Serratia.

### Analysis of electrochemical characteristics of plant tissue inoculation MFC

Voltage data was collected for MFC inoculated with angelica and sweet potato for 1500 hours.The MFC inoculated with angelica tissue gradually stabilized power generation after running for nearly 700 hours. The average power generation cycle reached 197.5 hours, and the average voltage reached 247.4 mV after stabilization. The MFC inoculated with sweet potato tissue started faster, and a stable power generation cycle began after 300 h, and the power generation cycle was shorter than that of Angelica tissue inoculation. The average power generation cycle was 185.7 h, which was stable. Afterwards, the average voltage reached 196.7 mV, which was lower than the MFC inoculated with Angelica sinensis.The voltage curve is shown in **Figure 3**.

**Figure 3.**
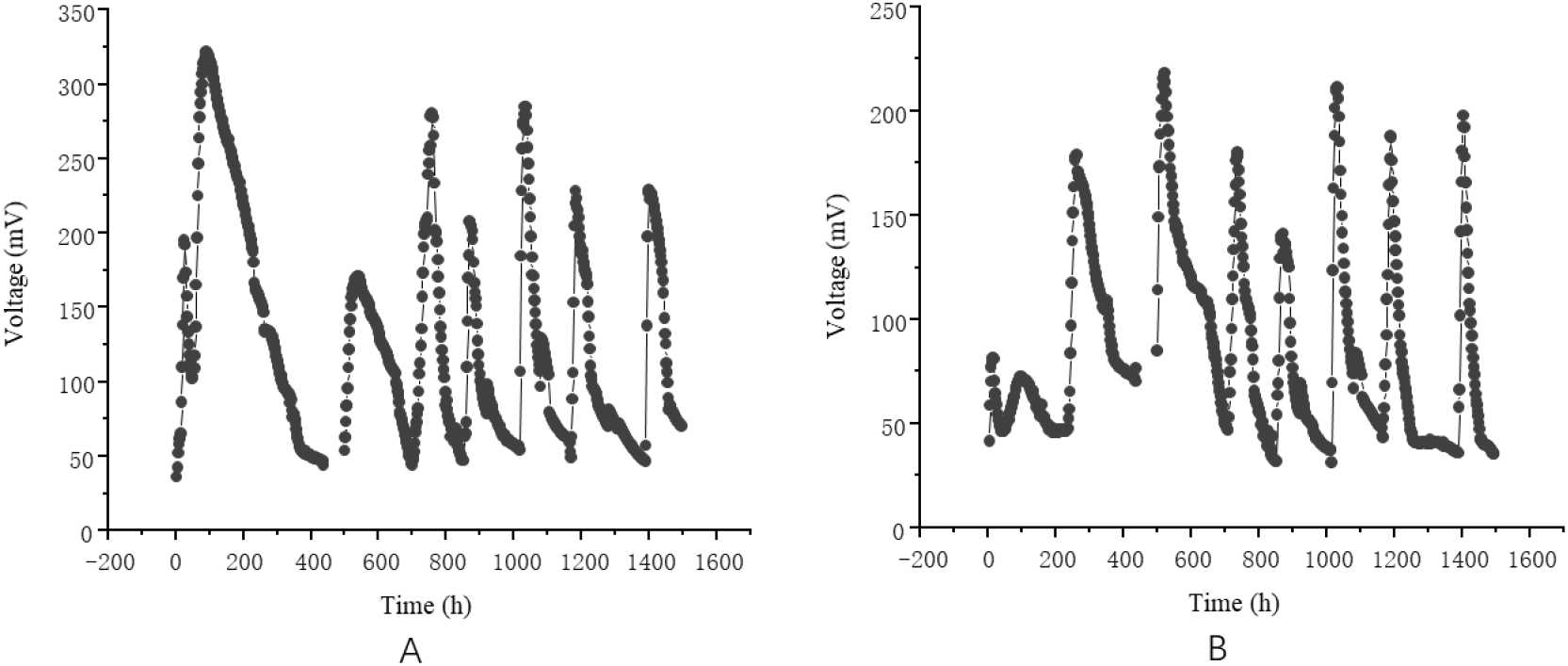
Figure 3-11 Voltage curve of angelica tissue (A) and sweet potato tissue (B) inoculated with MFC.

As shown in **Figure 4**, the MFC inoculated with angelica tissue showed a high power density, reaching 196.9±5.8 mW/m^2^, and the maximum power density of MFC inoculated with sweet potato tissue was relatively low, only reaching 142.9±16.4 mW/m^2^. Both the MFC inoculated with sweet potato and angelica produced maximum power density when the external resistance value was 1000 Ω, but the corresponding current density was different. Sweet potato reached 436.4 mA/m^2^ and Angelica reached 512.4 mA/m^2^. The voltage of MFC inoculated with angelica tissue dropped rapidly in the low current density and high current density areas, while the voltage of MFC inoculated with sweet potato tissue decreased slowly in the low current density area.

**Figure 4.**
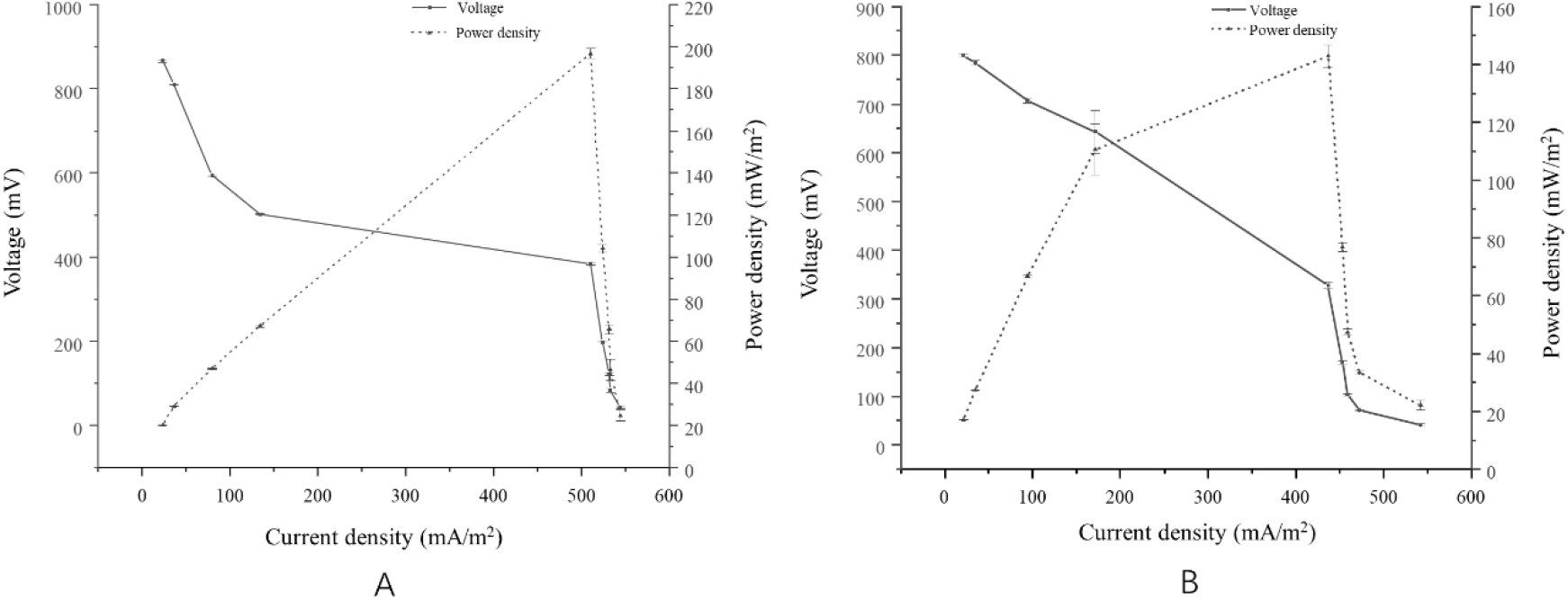
Power density curve and polarization curve of angelica tissue (A) and sweet potato tissue (B) inoculated with MFC

**Figure 5.**
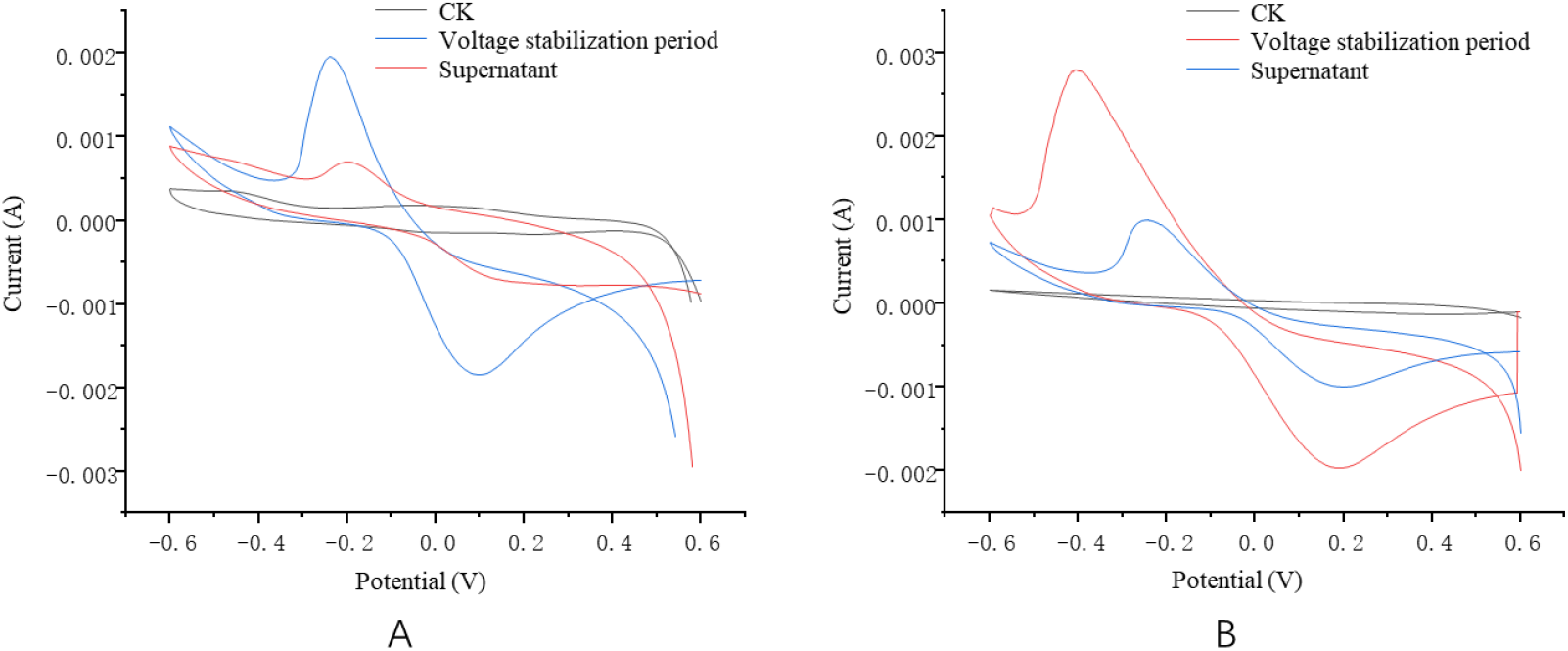
Cyclic voltammetry curve of angelic tissue (A) and a sweet potatotissue (B) inoculated with MFC The scanning electron microscope observation results of the MFC anode electrode inoculated with Angelica sinensis and sweet potato tissues are shown in **Figure 6**. The adhesion density of the bacterial cells on the carbon felt of Angelica anode electrode is large, and the bacillus occupies a large proportion in the field of view. The attachment density of the bacterial cells on the sweet potato anode electrode is relatively small, and the adhesion of bacillus and cocci can be observed.

As shown in Figure 3-15, the MFC in the stable period of Angelica sinensis tissue inoculation produced obvious redox peaks, which were located near −0.4 V and 0.2 V, respectively. The anode supernatant also showed good redox characteristics, respectively. Redox peaks appeared at −0.3 V and −0.2 V. Unlike Angelica sinensis, MFC inoculated with sweet potato had a slightly lower redox peak intensity at peak positions at −0.3 V and 0.1 V, respectively, and the supernatant exhibited a slightly lower redox peak than Angelica at −0.2 V and Appears at 0.1 V.

### Identification of strain H6 and analyses of its power production performance

#### Morphological characteristics

To verify the electricity production performance of endophytic bacteria, we selected a representative strain, H6, from the isolated strains. The strain H6 was assessed for its morphological characteristics after culturing on LB agar medium. The colonies of the isolate were cream-colored on NA, and the edge of the colonies was wavy. Moreover, the isolate was able to grow in 5%, 7%, 9%, 11%, 13% and 17% NaCl concentrations. The results of physiological and biochemical tests of the strain H6 are shown in **Table 2**.

**Table 2.** Results of physiological and biochemical tests of strain H6.

| Project | Result | Project | Result |
| --- | --- | --- | --- |
| Gram stain | + | 11% salinity | + |
| Glycolysis | + | 13% salinity | + |
| Amylase | + | 17% salinity | + |
| Gelatin liquefaction | + | Mannitol | + |
| Urease | + | Maltose | + |
| 5% salinity | + | Raffinose | - |
| 7% salinity | + | Lactose | + |
| 9% salinity | + | D- (+) - Trehalose dihydrate | + |

+: positive reaction; -: negative reaction

An approximately 1 500 bp fragment of the 16S rDNA gene region was also sequenced to further characterize the isolate and to construct a dendrogram using closely related species. Combined with physiological and biochemical identification results, the isolated was identified as *Staphylococcus sp*. These identifications were also supported by the phylogenetic analysis (**Figure 7**). Combined with physiological, biochemical and molecular biological identification, the strain H6 was a strain of *Staphylococcus succinus subsp. succinus*. The nucleotide sequence accession numbers and the GenBank database accession numbers for the 16S rDNA nucleotide sequences of the isolate are MK764983.

**Figure 6.**
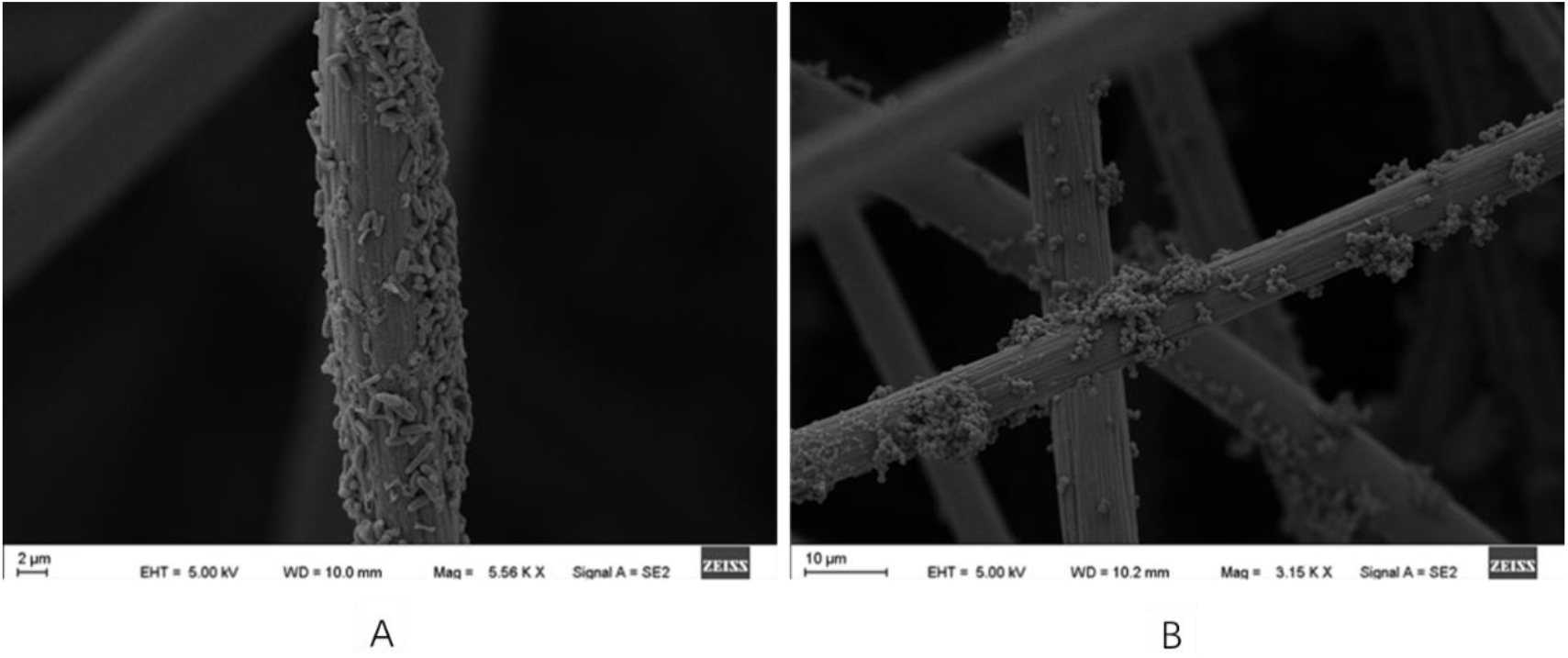
Scanning electron microscope of anode electrode of sweet potato tissue (a) and angelica tissue (b) inoculated with MFC

**Figure 7.**
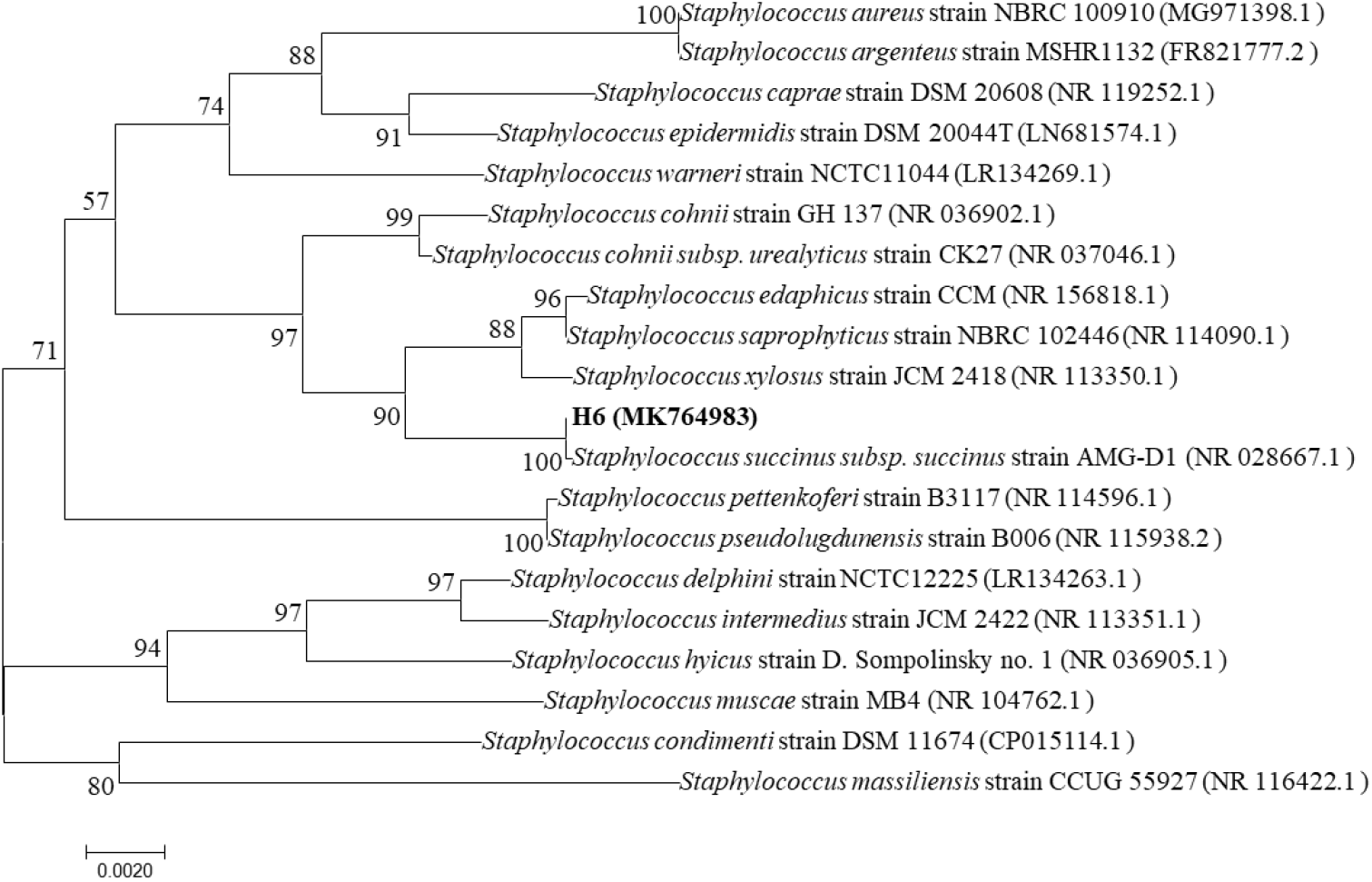
A neighbor-joining tree of the bacteria and their closely related species. The approximately 1 500-bp sequence of the 16S rDNA gene was used to construct a dendrogram. Bootstrap values based on 1 000 replicates are indicated above nodes. Bootstrap values C ≥ 50 are labeled. The scale at the bottom of the dendrogram indicates the degree of dissimilarity

After inoculation of H6, the cell voltage increased rapidly after a 20 h lag phase and became stable after the 150th hour. Each stable generation cycle was maintained for 75 hours. H6 produced a maximum voltage up to 130 mV with a 1 kΩ external resistance (**Figure 8**).

**Figure 8.**
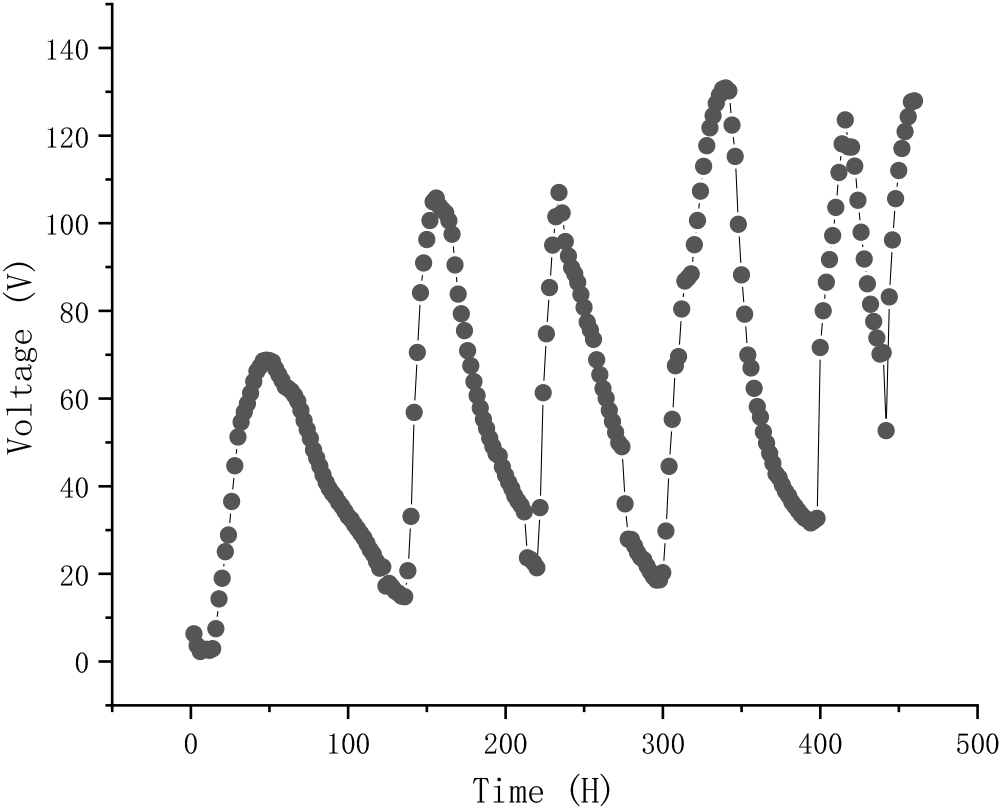
Voltage measured in the MFC that operated with a 1 kΩ external resistance.

Polarization and power density curves were obtained to determine the maximum currents and power densities (**Figure 9)**. The open circuit voltage was measured at an average of 742 mV. H6 produced a maximum power density of 1 910 mW/m^3^ at a resistance of 5 000 Ω, and the maximum current was 0.158 mA. The results indicated that H6 had significant electricity generation performance.

**Figure 9.**
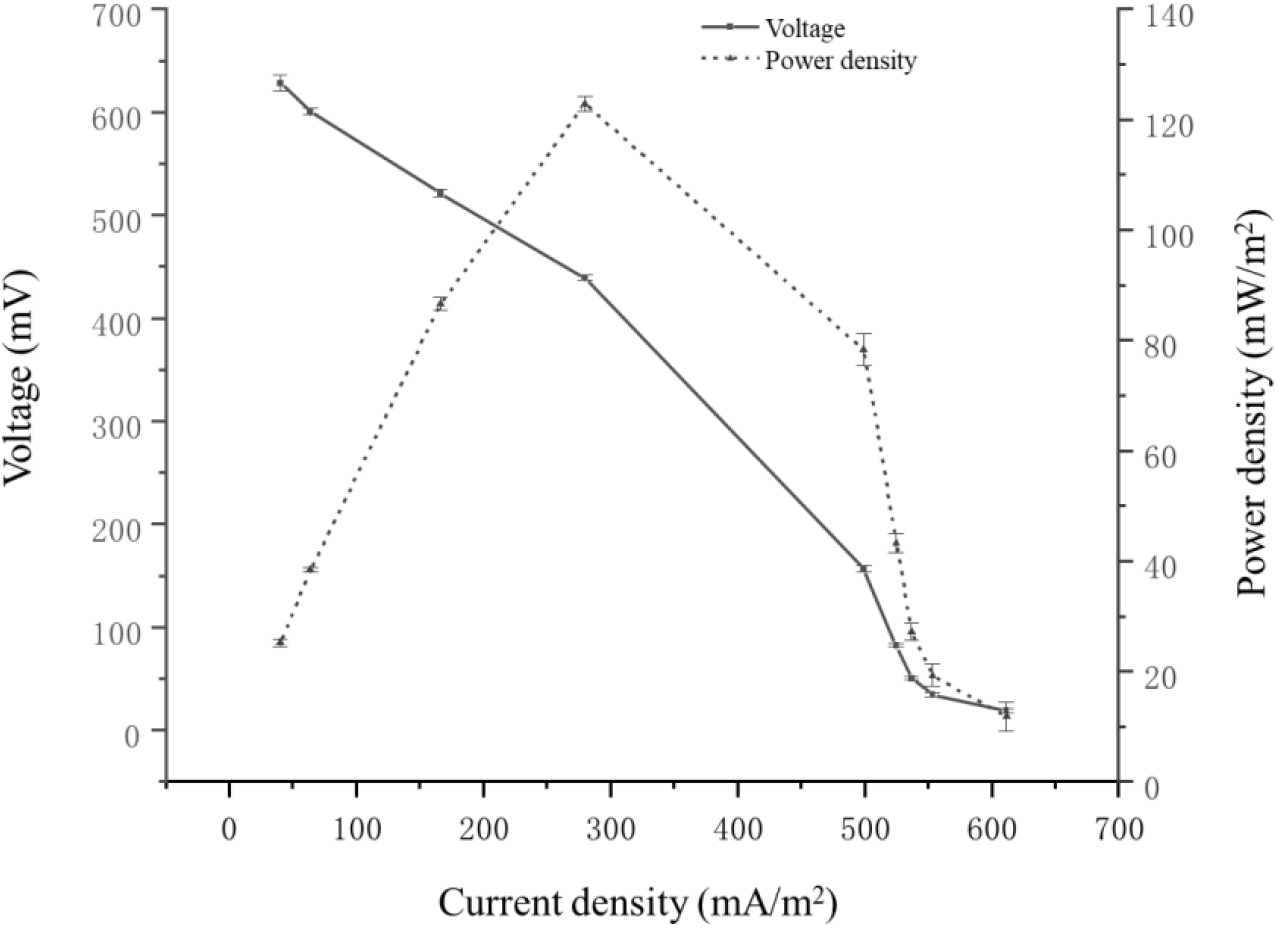
Polarization curves (lines) and power density curves (dotted line) for an MFC inoculated with H6. The test was conducted when the cell voltage reached the maximum and stabilized for several hours.

The fresh medium, voltage rise period of the MFC, voltage stabilization period of the MFC and the anode medium supernatant without suspended cells were investigated by CV to determine the electrochemical activity of H6 and self-secreted mediators. No electrochemical activity was observed when the fresh medium was tested. A voltammogram of the voltage rise period of the MFC showed an oxidization peak at −240 mV and a reduction peak at 200 mV (**Figure 10**). The voltage stabilization period of the MFC showed an oxidization peak at −210 mV (vs. SCE) and a reduction peak at 76 mV, and the anode medium supernatant showed an oxidization peak at −155 mV and a reduction peak at 140 mV. The results showed that their redox peaks were not at the same potential.

**Figure 10.**
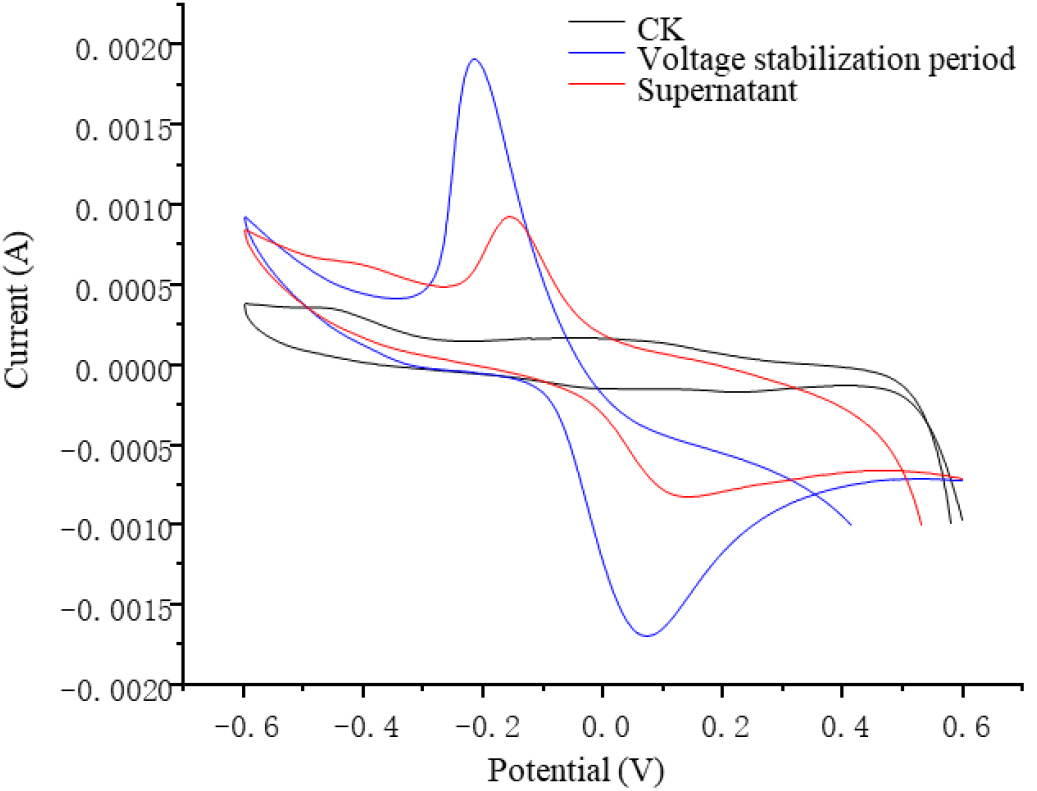
CV of an uninoculated MFC, voltage rise period of the MFC, voltage stabilization period of the MFC and centrifuged anode medium supernatant after the voltage stabilization period. The potential ranged from −600 mV to 600 mV, with a scan rate of 5 mV/s.

**Figure 11.**
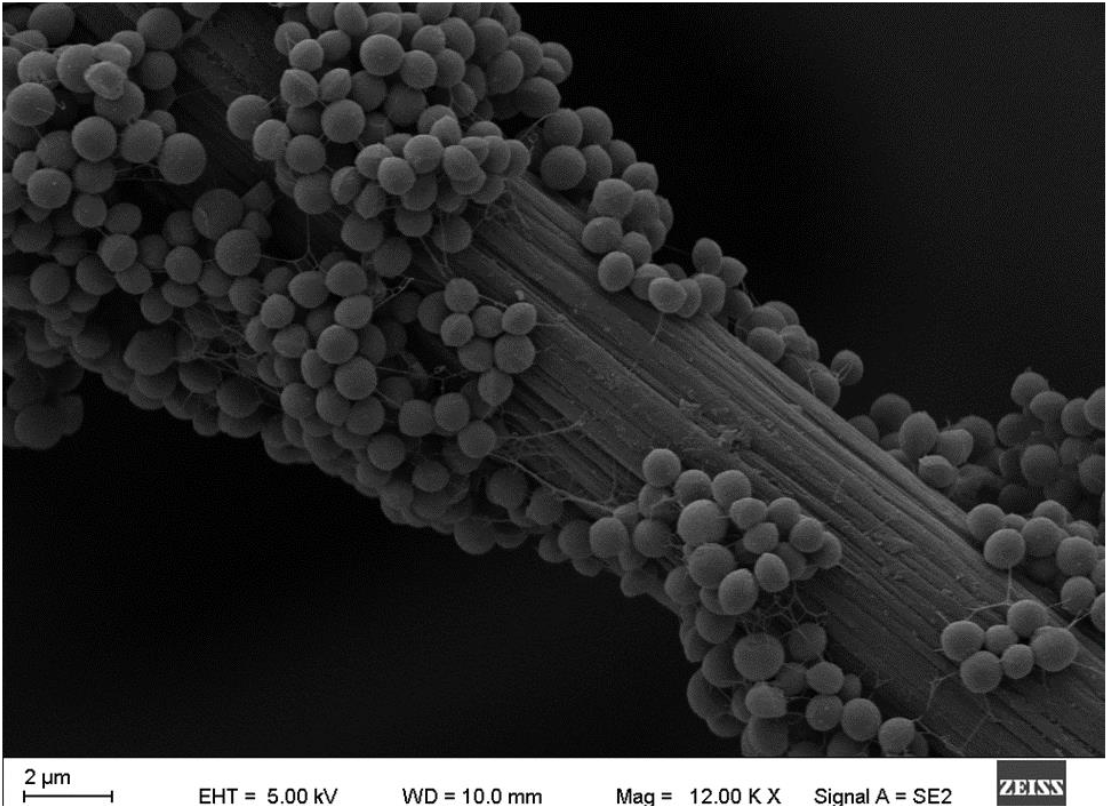
H6 inoculation MFC anode electrode scanning electron microscope

Observation by scanning electron microscope revealed that the spherical cells were densely attached to the carbon felt fibers. It is particularly worth noting that the strain produced filaments in contact with the electrode, and its main components and whether they play the role of nanowires still need further study.

## Discussion

Microorganisms that can generate an electrical current have been of scientific interest for more than one century. Electroactive microorganisms as anode catalysts have a vital impact on the application of MFCs. Electroactive microorganism resources are also widely explored^(24)^. Because MFCs are widely used in wastewater treatment, electroactive microorganism resources are usually isolated with all kinds of sewage ^(9, 25, 26)^. Although most of the studies on the resources of electroactive microorganisms have been concentrated in wastewater, it is undeniable that the electroactive microorganisms are widely distributed. In recent years, researchers have isolated many strains of electroactive microorganisms from various ecological environments. Many different types of environments harbor electroactive microorganisms, including anaerobic sludge from treatment plants, anaerobic sediment, and even soil^(27)^. However, thus far, no studies have reported the existence of electroactive microorganisms in plant endophytic bacteria resources.

In our study, we isolated 26 endophytic bacteria belonging to 10 genera from sweet potato and angelica. Through further screening, 17 strains belonging to 6 genera were shown to have electrochemical activity. Because of the difference in living environment and metabolites between the two plants, there were significant differences in their endophytic bacteria resources. The diversity of endophytic bacteria in angelica was richer than that in sweet potato, but the number of endophytic bacteria was lower in angelica than in sweet potato. Plant endophytic bacteria show diversity as potential resources of electroactive microorganisms. These electroactive microorganisms are concentrated in mainly *Staphylococcus sp*., *Shinella sp*., *Pleomorphomonas sp*., *Paenibacillus sp*., and *Bacillus sp*. genera. This distribution is very different from the species distribution of electroactive microorganisms isolated in the past. So far, most of the electroactive microorganisms found are concentrated in *Shewanella sp*., *Geobacter sp*., *Pseudomonas sp*., *Klebsiella sp*. and so on ^(24)^. In general, *Pseudomonas sp*. are considered to have good extracellular electron transport ability ^(28, 29)^, but only two of the 6 endophytic strains isolated from plants had electrochemical activity. This result indicates that *Pseudomonas sp*. in endophytic bacteria is not the dominant species of electrogenic bacteria. The strain belonging to *Paenibacillus sp*. was the only one found with electrochemical activity from the tissues of sweet potato. The electrogenic activity of bacteria from *Pleomorphomonas sp*., *Rahnella sp*., *Shinella sp*. and *Staphylococcus sp*. was discovered for the first time. In particular, the isolates of *Staphylococcus sp*. were abundant. All of those strains have been widely isolated from plants, with a wealth of diversity^(30-34)^. Moreover, it has been reported that these species of bacteria have good pollutant degradation ability and no pathogenicity, so they can be used widely in a variety of MFCs ^(32, 35-37)^.

Some studies have shown that plant endophytic bacteria resources are very rich and have a variety of biological activities ^(38, 39)^ (in this study, as many as 26 endophytic bacteria belonging to 10 genera were isolated with partially culturable endophytic bacteria from only two plants by pure culture, and considering that the diversity of endophytic bacteria in plants was much more than that, it can be concluded that plant endophytic bacteria resources are a huge treasure trove for screening electroactive microorganisms.

Compared with pure culture, more electricity-producing microorganisms can be enriched by using MFC, especially some non-cultivable microorganisms. The high-throughput sequencing method can be used to analyze the microbial community structure in the MFC anolyte and the electrode membrane in the stable period, and then understand the potential of electricity-producing microorganisms. In the experiment, the anode microbial community of MFC stably inoculated with two plant tissues was analyzed. There is no single dominant population in the bacterial community developed on the anode, which may be because there is more than one type of electricity-producing microorganism in the system. The diversity of bacterial communities attached to the MFC electrode membrane inoculated with Angelica sinensis was lower than that of the anolyte, and the sequencing results of sweet potato showed that more and more abundant bacteria were attached to the electrode membrane. Proteobacteria bacteria account for a large proportion in the MFC anode solution and anode electrode membrane inoculated with Angelica sinensis, and the MFC anode electrode community inoculated with sweet potatoes is also dominated by Proteobacteria bacteria. However, the sweet potato anolyte is completely different. Bacteria in the rear wall bacteria occupy almost the majority of the community in the MFC anolyte. Most of the MFC anode community structure is dominated by Proteobacteria, and the current isolated and identified electro-producing bacteria are also concentrated here. For example, in the multiple groups of wetland plant-sediment MFCs built by Zhu Juanping, the dominant bacteria are all Proteobacteria ^(40)^; Chae et al. inoculated 4 MFCs with anaerobic sludge, and conducted feeding experiments on 4 different substrates for more than 1 year, and abundant proteobacteria were observed in all MFCs Bacteria ^(41)^. Among the MFCs inoculated with Angelica sinensis, the predominant bacteria in AC and AM were Pseudomonas, Lachnoclostridium 5, Clostridioides, Clostridium, Bacillus pumilus, and Clostridium spp. 1 as well as MFC inoculated with sweet potatoes. The dominant genus Staphylococcus and Lachnoclostridium 5 are not common in the MFC anodes of other inoculums^(42)^, while the common genus Geobacter, Klebsiella, and Bacteroide have not been identified in both MFCs. This difference in MFC inoculated with Angelica sinensis and sweet potato proves that the community structure of the electricity-generating microorganisms in plant tissues is completely different from other environments. Plant endophytes have great potential and advantages as a resource bank for electricity-producing microorganisms. The microbial community structure in a wide range of plant tissues is diverse, and the completely different community structure also provides certain guidance for the combined application of electricity-producing microorganisms.

To further verify the electricity production performance of endophytic bacteria, we selected the strain H6 from the dominant *Staphylococcus sp*. to be tested. The strain H6 was identified as *Staphylococcus succinus subsp. succinus* by morphology, physiology, biochemistry and molecular biology. It was found that the strain H6 had strong salinity and high pH tolerance. *Staphylococcus succinus subsp. succinus* is a gram-positive coccoid bacterium first identified in 1998 after its isolation from Dominican amber ^(43)^. This species have since been isolated from diverse environments, including cheese, dry or fermented meat products, human clinical specimens, the Dead Sea, plants, soil and fragments ^(44-48)^. Julianne and Brendan identified several genes that were associated with resistance to heavy metals and toxic compounds (copper, cobalt, zinc, cadmium, mercury, arsenic, and chromium). Therefore, the strain H6 has good application prospects in wastewater treatment.

The cell was loaded with an external resistance of 1 000 Ω, and the change in current was recorded. We observed five cycles of stable voltage generation in 470 hours. After replacing the anode medium in each cycle, it showed the ability to rapidly recover the voltage. The strain H6 showed good power production ability.

In this study, we showed evidence for direct and indirect electron flow from H6 to the electrode using the cyclic voltammetry technique and the fuel cell. With the progress of battery power generation, the redox peak of the cyclic voltammetry curve in the voltage stability period was significantly higher than that in the voltage rise period. However, the cyclic voltammetry curve of the supernatant after centrifugation still has a considerable redox peak, which indicates that the bacteria produced electron intermediaries to transfer electrons^(49)^. In additionthe redox peak of the supernatant cyclic voltammetry curve was significantly weak, and the voltage was stable, indicating that H6 may have other ways of transferring electrons to the electrode, such as through direct contact with the electrode ^(50, 51)^. We will further study the H6-specific electron transfer mechanism at a later stage. Electroactive microorganisms, which can transfer electrons by self-secreted electron intermediaries, have been widely studied^(38, 52)^. In practical applications, the efficiency of MFCs is greatly improved.

The MFC inoculated with Angelica sinensis and sweet potato tissue showed a higher power density than single bacteria, reflecting the advantages of mixed endophyte mixed community power generation. Studies have shown that the power generation capacity of the MFC system constructed by pure microbial culture and the adaptability to complex environments are lower than the MFC system constructed by mixed bacteria (53), but pure culture helps to clarify the electron transfer mechanism at the microbial level, And further reduce the complexity of mixed communities. Future research directions should focus on the screening, domestication, modification and optimization of multi-species to improve their electrochemical activity.

Why plant endophytic strains have electricity-generating properties, and what is the significance of this special ability? These are some important questions. Several isolated plant-producing electric bacteria have been reported to have a certain nitrogen-fixing effect. The analysis suggests that the plant-producing electric bacteria may have an important interaction relationship with their endophytic or rhizosphere nitrogen-fixing bacteria. Most of the heterotrophic bacteria found in natural soil currently depend on nitrogen sources (nitrate or ammonium) from free-living or symbiotic nitrogen-fixing bacteria, but there are also some that can metabolize and fix nitrogen in the atmosphere under limited nitrogen conditions(54). Studies have shown that the high balance of NADH and ATP in the metabolism of nitrogen-fixing bacteria may be caused by the electrochemical reduction force, and they are also factors which activate nitrogen fixation(55). Under the condition of limited organic carbon, electrochemically induced nitrogen fixation may be an important way to increase the nitrogen source (56). Jung et al. built a conventional bioreactor and an electrochemical bioreactor, and evaluated the effect of using autogenous nitrogen-fixing bacteria to biochemically convert nitrogen in the atmosphere into ammonia nitrogen using electrochemical reduction. The biomass and ammonia nitrogen content in the bacterial culture discharged from the electrochemical bioreactor are 1.6 times and 6 times that of the conventional bioreactor, respectively, and the analysis suggests that the electrochemical reducing power may activate the nitrogen-fixing effect of nitrogen-fixing bacteria (57). Taking into account that the extracellular electron transport capacity of electro-producing bacteria in plants may also provide a certain electrochemical reduction force for the nitrogen-fixing bacteria’s nitrogen-fixing effect. Whether there are other interactions between the endophytic bacteria and other plant endophytes and their rhizosphere nitrogen-fixing bacteria remains to be further studied.

The present research provided a preliminary understanding of the diversity of electroactive microorganisms in plants and showed a different resource pool of electroactive microorganisms from those of previous studies. Why there are so many electroactive microorganisms in plants remains a question, and we will continue to research and think about it in the future. Moreover, whether there is a functional relationship between plants and the extracellular electron transfer of their endophytic bacteria also needs to be further studied.

## Acknowledgements

This work was supported in part by Lanzhou Science and Technology Plan Project 2018-1-104; Northwest Normal University Innovation Capacity Improvement Program CXCY2018B009; Gansu International Science and Technology Cooperation Special 1504WKCA028.

## Declarations

### Ethics approval

This article does not contain any studies with human participants or animals performed by any of the authors.

### Consent to participate

All authors clarify and agree to participate in relevant research.

### Consent for publication

All authors clarify and agree to publish this manuscript

### Availability of data and material

All test data in the article are transparent and materials are available.

### Code availability

Not applicable

### Authors’ contributions

LL conceived and designed research. ZL and YZ conducted experiments. CY and WM analyzed data. ZL and LL wrote the manuscript. All authors read and approved the manuscript.

---

**Figure 2** A neighbor-joining tree of the bacteria and their closely related species. The approximately 1 500-bp sequence of the 16S rDNA gene was used to construct a dendrogram. Bootstrap values based on 1 000 replicates are indicated above nodes. Bootstrap values C ≥ 50 are labeled. The scale at the bottom of the dendrogram indicates the degree of dissimilarity

**Figure 3** Voltage measured in the MFC that operated with a 1 kΩ external resistance.

**Figure 4** Polarization curves (lines) and power density curves (dotted line) for an MFC inoculated with H6. The test was conducted when the cell voltage reached the maximum and stabilized for several hours.

**Figure 5** CV of an uninoculated MFC, voltage rise period of the MFC, voltage stabilization period of the MFC and centrifuged anode medium supernatant after the voltage stabilization period. The potential ranged from −600 mV to 600 mV, with a scan rate of 5 mV/s.

